# Generation and characterization of iPSC lines (UOHi001-A, UOHi002-A) From a patient with SHANK3 mutation and her healthy mother

**DOI:** 10.1101/2022.04.13.486968

**Authors:** Ritu Nayak, Idan Rosh, Tatiana Rabinski, Menachem Mendel Percia, Shani Stern

## Abstract

Phelan-McDermid syndrome (PMS) is a rare genetic condition that causes global developmental disability, delayed or absent speech, and autism spectrum disorder. The loss of function of one copy of *SHANK3*, which codes for a scaffolding protein found in the postsynaptic density of synapses, has been identified as the main cause of PMS. We report the generation and characterization of two induced pluripotent stem cell (iPSC) lines derived from one patient with a *SHANK3* mutation and the patient’s mother as a control. Both lines expressed pluripotency markers, differentiated into the three germ layers, retained the disease-causing mutation, and displayed normal karyotypes.

## Resource Table

**Resource Table:**
|  |  |
| --- | --- |
| Unique stem cell lines identifier | 1. UOH001-A<br>2. UOH002-A |
| Alternative name(s) of stem cell lines | UOH001iShank3<br>UOH002iShank3 |
| Institution | University of Haifa (UOH), Ben Gurion University of the Negev (BGU) |
| Contact information of distributor | Shani Stern |
| Type of cell lines | iPSC |
| Origin | Human |
| Additional origin info required | UOH001iShank3 : 10 years old, sex- Female, Caucasian<br>UOH002iShank3: 47 years old , sex-Female, Caucasian |
| Cell Source | Skin fibroblasts |
| Clonality | Clonal |
| Associated disease | Phelan-McDermid Syndrome |
| Gene/locus | SHANK3 |
| Date archived/stock date | 22<sup>nd</sup> March, 2022 |
| Cell line repository/bank | https://hpscreg.eu/cell-line/UOH001-A |
| Ethical approval | The Sheba Medical Centre Institutional Helsinki committee<br>Approval number: SMC-6158-19 |

## Resource utility

In the future, the generated patient and unaffected mother iPSC lines will aid in the development of a disease-in a dish model for PMS. To investigate the neuronal function and molecular mechanisms associated with the *SHANK3* (C.3679insG) mutation, iPSCs can be differentiated into *SHANK3*-mutant neurons of the cortex or hippocampus and their functional and cellular alterations can be measured similar to measurements in other ASD patients derived neurons (1, 2).

## Resource Details

Phelan-Mcdermid syndrome (PMS), is caused by a mutation in the *SHANK3* gene. It is a congenital condition that can affect people of all genders (3). Literature survey suggests that the phenotypic expression of PMS can involve one or more haploinsufficent gene. *SHANK3* gene has emerged as a robust candidate gene in neurobehavioral aspects of PMS (3, 4).The neurobehavioral symptoms is characterised by a developmental delay, a moderate to an extreme intellectual impairment, low muscular tone or hypotonia, and delayed speech. It is estimated that more than 80% of those with PMS meet the diagnostic criteria for autism or autistic-like behaviour(5).

Here, we report the details regarding the generation and characterization of two iPSC lines derived from a 10-year-old female patient with *SHANK*3 mutation, and her unaffected mother (age 47). Fibroblasts were harvested from skin biopsies and collected from both the candidates and were grown on six well plates (Fig. 1A). Using electroporation, fibroblasts were reprogrammed to generate iPSCs (Fig. 1B) using non-integrating episomal plasmids encoding human OCT3/4, SOX2, KLF4, L-Myc, shp53, Lin28 and SV40LT as previously described (Fig. 1C) (Vatine et al., 2017).Immunocytochemistry (ICC) was performed to confirm the expression of pluripotency markers in the two lines. TRA-1-60, SSEA-4, OCT 3/4, SOX2, and NANOG were all expressed by both lines. Fluorescence-activated cell sorting (FACS) analysis revealed that pluripotency markers such as TRA-1-60, SSEA-4, OCT3/4, SOX2 and NANOG were expressed in atleast 85% of the cells. All iPSC clones generated embryoid bodies (EBs) and spontaneously differentiated into the three germ layers. These germ layers were validated by immunocytochemistry (ICC) for three-germ layer specific markers such as neurofilament, SMA and fetoprotein. All lines had a normal karyotype and were clear of mycoplasma.

**Fig. 1.**
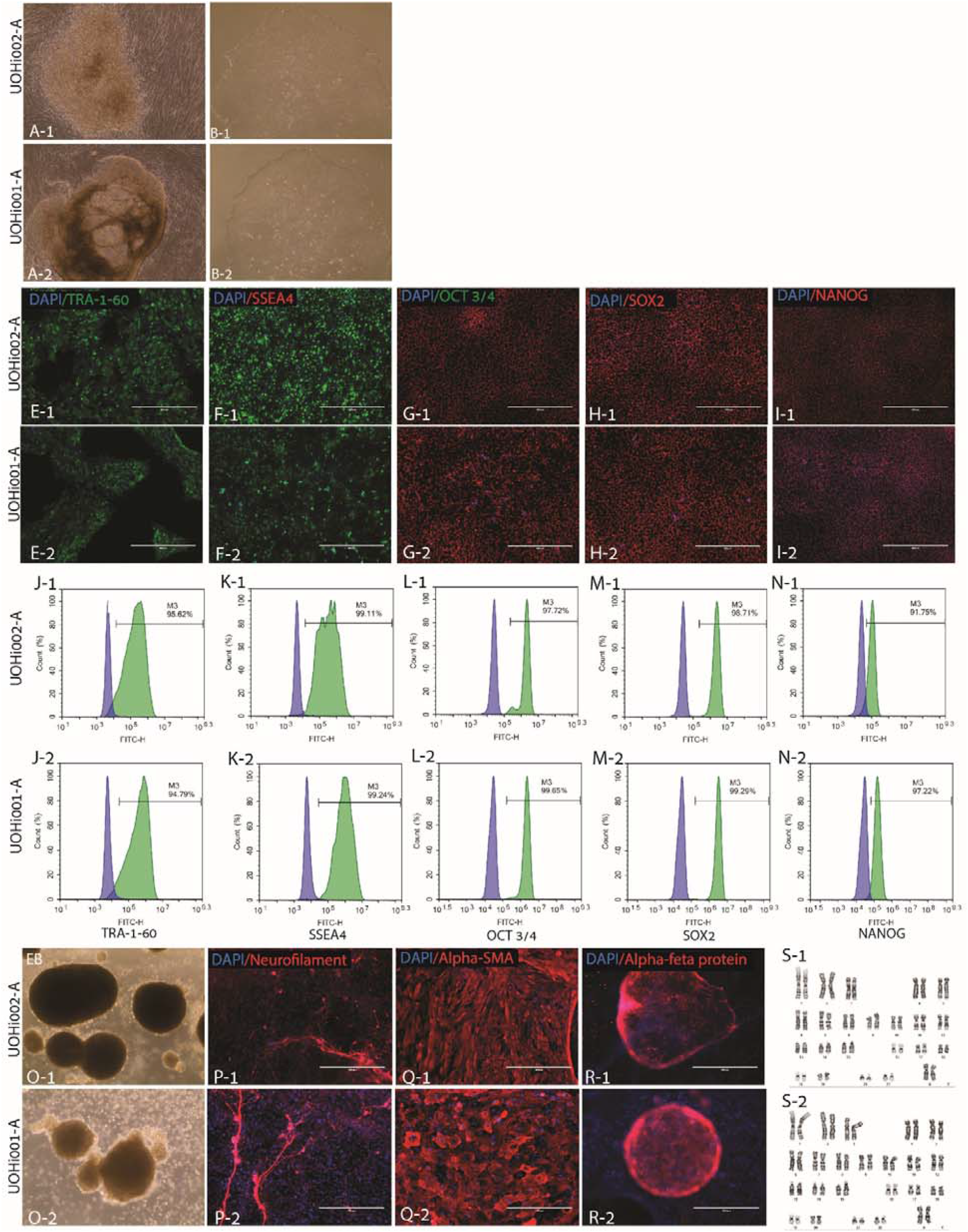
Characterization of UOHi001-A and UOHi002-A iPSCs.

## Materials and Methods

### 1. Cell culture conditions

All cells were grown in an incubator at 37 °C and 5% CO2.

### 2. Generation of patient-specific fibroblast

Skin biopsies were collected, dissected and grown in 6-well plates with coverslips for two weeks in DMEM (Biological industries, BI) with 20% fetal bovine serum (FBS) and 50% medium replacement every two days.

### 3. Reprogramming

A total of 10^6^ cells were collected with TrypLE (Gibco) and electroporated with non-integrating episomal vectors using a Neon transfection system (Invitrogen). Cells were cultured in DMEM media using 15% FBS, 5ng/ml basic fibroblast growth factor (bFGF, peprotech), and 5µM rock inhibitor on mouse embryonic fibroblast (MEF)-coated plates (Sigma-Aldrich).After 2 days, Nutristem media (BI) supplemented with 5ng/ml bFGF was utilized with medium replaced every other day. Six colonies were transferred to MEF-coated plates with Nutristem media containing 5ng/ml bFGF. Out of six, three colonies were selected and manually transferred to matrigel (corning) -coated plates with Nutristem, which was replenished daily.

### 4. Immunocytochemistry

iPSCs were rinsed twice with Dulbecco’s Phosphate-Buffered Saline (DPBS) and then fixed in 4% paraformaldehyde for 20 minutes at room temperature (RT), and washed again with DPBS. The fixed cells were treated with a blocking solution comprising of 1% bovine serum albumin and 0.1% triton-X-100 for one hour, followed by an overnight incubation with a primary antibody in blocking solution at 4°C. Cells were washed twice with blocking solution before being treated with fluorescently labelled secondary antibodies overnight at 4°C. The cell nuclei were stained with DAPI.

### 5. FACS analysis

Using TrypLE, iPSCs were dissociated and collected into single cells. Cells were rinsed with 3% FBS in DPBS before being incubated with a primary antibody solution at room temperature for 2 hours. For intracellular marker staining, cells were fixed for 40 minutes at room temperature using fixation solution (Invitrogen), rinsed with permeabilization solution (Invitrogen), and resuspended in primary antibody solution for two hours at room temperature. Cells were resuspended in 3% FBS in DPBS with primary antibodies and incubated for 40 minutes at 4⍰C for surface markers. Then, the cells were washed twice with DPBS and incubated in permeabilization solution containing secondary antibodies for intracellular markers and DPBS containing 3% FBS for surface markers. A NovoCyte flow cytometer was used for the analysis.

### 6. Differentiation potential

When the cells were confluent, iPSCs were collected and dissociated into single cells using TrypLE. Thereafter, the cells were resuspended in Nutristem media and supplemented with 10ng/ml bFGF and 7µM of ROCK inhibitor (Enzo life sciences) until EBs appeared spontaneously. The suspended EBs were fed with EB media containing 15% FBS, 1% Non-Essential Amino Acids, NEAA, and 0.1mM β-mercaptoethanol (Biological Industries). After 4-7 days, EBs were plated on 0.1% gelatin-coated plates and cultured for 21 days with the EB medium replaced twice a week. The cells were fixed and stained on day 21.

### 7. karyotyping

iPSCs were treated with 100ng/ml colcemid (BI), incubated for 60 minutes, and collected in versene (Gibco). Cells were preserved in a 1:3 glacial acetic acid: methanol (Biolabs-chemicals) solution and karyotyped using g-banding.

### 8. STR analysis

Profiling was performed for fibroblasts and derived iPSCs. These loci were tested: D8S1179, D21S11, D7S820, CSF1PO, D3S1358, TH01,D13S317, D16S539, D2S1338, D19S433, vWA, TPOX, D18S51, Amelogenin, D5S818, FGA.

### 9. Mycoplasma

Using the Hymycoplasma PCR kit, all lines were screened for mycoplasma contamination (Hylabs).

## Discussion

The reprogrammed LCLs will be further utilized as a useful cell line model that retains the patients’ genetic background and phenotypes including molecular and cellular properties such as gene regulatory pathways and functional alterations. The LCL-derived induced pluripotent stem cells will be differentiated into human-derived neurons and brain organoids. Using electrophysiological and molecular techniques, specific ionic channels or other neurophysiological changes can be detected. We expect to find functional differences since the Shank3 gene was shown to have an important role in the formation of efficient synapses (6). These measurements will aid in the understanding of disease mechanisms and novel drug discovery.

## Additional files

**Table 1:** Characterization and validation.

| Classification | Test | Result | Data |
| --- | --- | --- | --- |
| **Morphology** | Photography Bright field | Normal | Figure 1A and B |
| **Phenotype** | Qualitative analysis by Immunocytochemistry | Expression of Oct3/4, SSEA4, TRA-1-60, SOX2 and Nanog | Figure 1E-I |
|  | Quantitative analysis by Flow Cytometry | TRA-1-60: 95.62%, 94.79%<br>SSEA4: 99.11%, 99.24%<br>Oct 3/4: 97.72%, 99.65%<br>SOX2: 98.71%, 99.29%<br>Nanog: 91.75%, 97.22% | Figure 1J-N |
| **Genotype** | Karyotype (G-banding) and resolution | 46XX<br>Resolution 400 | Figure 1S |
| **Identity** | Microsatellite PCR (mPCR) OR | Not performed | *e.g. supplementary file 2* |
|  | STR analysis | Unknown | *e.g. submitted in archive with journal* |
| **Mutation analysis (IF APPLICABLE)** | Sequencing | Unknown | *e.g. Figure 1 panel D* |
|  | Southern Blot OR WGS | N/A | N/A |
| **Microbiology and virology** | Mycoplasma detection by PCR | Negative | Supplementary |
| **Differentiation potential** | Embryoid body formation and spontaneous differentiation | Expression of layer specific markers: 68 kDa Neurofilament (Ectoderm), α-Fetoprotein (endoderm), α-smooth muscle actin (mesoderm). | Figure 1O-R |
| **List of recommended germ layer markers** | **Expression of these markers has to be demonstrated at mRNA (RT PCR) or protein (IF) levels, at least 2 markers need to be shown per germ layer** |  |  |
| **Donor screening (OPTIONAL)** | HIV 1 + 2 Hepatitis B, Hepatitis C | N/A | *e.g. not shown but available with author* |
| **Genotype additional info (OPTIONAL)** | Blood group genotyping | N/A | *e.g. not shown but available with author* |
|  | HLA tissue typing | N/A | *e.g. not shown but available with author* |

**Table 2:** Reagents details.

|  | **Antibodies used for immunocytochemistry/flow-cytometry** |  |  |  |
| --- | --- | --- | --- | --- |
|  | **Antibody** | **Dilution** | **Company Cat #** | **RRID** |
| Pluripotency Markers | Mouse anti OCT3/4 | 1:100 | Santa Cruz, sc-5279 | RRID: AB_628051 |
|  | Mouse anti TRA-1-60 | 1:50 | R&D, MAB4770 | RRID: AB_2119062 |
|  | Mouse anti SSEA-4 | 1:100 ICC | Santa Cruz, SC-21704 | RRID:AB_628289 |
|  | Mouse anti IgG3 | 1:100 FACS |  |  |
|  | Isotype | 1:100 | Bio-legend, 330405 | RRID: AB_1089207 |
|  | Rabbit anti SOX2 | 1:100 | Abcam, ab97959 | RRID: AB_2341193 |
|  | Rabbit anti NANOG | 1:100 | Abcam, ab21624 | RRID: AB_446437 |
| Differentiation Markers | Rabbit anti 68kDa Neurofilament | 1:100 | Abcam, ab52989 | RRID: AB_869924 |
| | Rabbit anti α-Fetoprotein | 1:1 | ScyTek, A00058 | RRID:AB_651223 |
| | Rabbit anti α-smooth muscle actin | 1:100 | Abcam, ab32575 | RRID: AB_722538 |
| Secondary antibodies | Alexa Fluor 488-conjugated Anti Mouse | 1:500 | Jackson ImmunoResearch inc., 715-545-150 | RRID: AB_2340846 |
|  | Alexa Fluor 594-conjugated Anti Rabbit | 1:500 | Jackson ImmunoResearch inc., 715-545-152 | RRID: AB_2340621 |
|  | Alexa Fluor 488-conjugated Anti Rabbit | 1:500 | Molecular Probes, A-11029 | RRID:AB_138404 |
|  | Alexa Fluor 488-conjugated Anti Mouse | 1:500 | Thermo Fisher Scientific, A-11034 | AB_2576217 |

