## Supplementary material for "Generation and characterization of iPSC lines (UOHi001-A, UOHi002-A) From a patient with SHANK3 mutation and her healthy mother": Supp 1

| Date of receiving: | 16.08.2021 | Hylabs test: | 66-81 |
|---|---|---|---|
| Company: | BGU | Order ID: | 395825 |
| Name: | Meshi Zorsky | Version No: | 1 |
| E-mail: | | | |
| Mobile: | | | |

### 1. Samples:

| number | **Sample** | number | **Sample** | **number** | **Sample** |
|---|---|---|---|---|---|
| 1 | 001Shank3Fbp3 | 9 | 009HikeshiCTRFbp2 | 17 | 011PTHSCTRFbp2 |
| 2 | BGU001iShank3n6p5 | 10 | BGU009iHikeshiCTRn2p8 | 18 | BGU011iPTHSCTRn5p21 |
| 3 | 007Shank3CTRFbp2 | 11 | 001STXBP1Fbp1 | 19 | 012PTHSCTRFbp3 |
| 4 | BGU002iShank3CTRn1p5 | 12 | BGU001iSTXBP1n6p13 | 20 | BGU012iPTHSCTRn6p15 |
| 5 | 009HikeshiFbp2 | 13 | 002STXBP1CTRFbp2 | 21 | 013PTHSFbp2 |
| 6 | BGU009iHikeshin1p7 | 14 | BGU002iSTXBP1CTRn2p9 | 22 | BGU013iPTHSn3p20 |
| 7 | 010HikeshiFbp2 | 15 | 011PTHSFbp3 | 23 | 013PTHSCTRFbp3 |
| 8 | BGU010iHikeshin4p7 | 16 | BGU011iPTHSn3p21 | 24 | BGU013iPTHSCTRn4p22 |

### 2. Related Documents

2.1 Hy Laboratories SOP No. 11-032- "Operating Instructions for Nano Drop Instrument".

2.2 Hy Laboratories SOP No. 11-003- "DNA Fragment Analysis by Capillary Electrophoresis (STR)".

2.3 Hy Laboratories SOP No. 11-003- "Sequencing of DNA with Genetic Analyzer"

### 3. Materials and Software

3.1 AmpFLSTR™ Identifiler™ Plus PCR Amplification Kit (Thermos Fisher, Cat: 4427368)

3.2 Mapmarker Dy™ 632 50-500 (Hylabs)

3.3 Gene Mapper software v.6 (Thermo fisher)

### 4. Results:

4.1 Tables below summarized the results obtained for each cell line in the listed markers.

4.2 Raw date for each sample presented in ZIP file.

This report is submitted for the exclusive use of the person or corporation to whom is addressed, and neither the report nor the name of these laboratories nor any members of its staff may be used in connection with the advertising, sale or any product or process, or any non-regulatory third party without the written permission of Hy-Laboratories Ltd. The results are approved for samples tested only. For research use only.

| Locus designation | 001Shank3Fbp3 | BGU 001iShank3n6p5 | 007Shank 3CTRFbp2 | BGU 002iShank3CTRn1p5 | 009Hikeshi Fbp2 | BGU 009iHikeshin1p7 |
|---|---|---|---|---|---|---|
| D8S1179 | 15 | 15 | 14, 15 | 14, 15 | 10, 14 | |
| D21S11 | 29 | 29 | 28, 29 | 28, 29 | 30, 32 | |
| D7S820 | 10 | 10 | 10, 13 | 10, 13 | 10, 11 | |
| CSF1PO | 10, 12 | 10, 12 | 12 | 12 | 8, 12 | |
| D3S1358 | 15, 18 | 15, 18 | 16, 18 | 16, 18 | 14 | |
| TH01 | 7, 10 | 7, 10 | 9.3, 10 | 9.3, 10 | 8, 9.3 | |
| D13S317 | 9, 11 | 9, 11 | 9, 12 | 9, 12 | 11, 12 | |
| D16S539 | 11 | 11 | 11 | 11 | 8, 11 | |
| D2S1338 | 17, 20 | 17, 20 | 19, 20 | 19, 20 | 23, 25 | |
| D19S433 | 13 | 13 | 13, 15.2 | 13, 15.2 | 12, 13 | |
| vWA | 16, 18 | 16, 18 | 16, 18 | 16, 18 | 16, 18 | |
| TPOX | 9, 11 | 9, 11 | 9, 11 | 9, 11 | 8, 9 | |
| D18S51 | 12 | 12 | 12 | 12 | 15, 20 | |
| AMEL | X | X | X | X | X | |
| D5S818 | 9, 11 | 9, 11 | 9, 10 | 9, 10 | 11, 13 | |
| FGA | 20, 26 | 20, 26 | 26, 27 | 26, 27 | 21 | |

This report is submitted for the exclusive use of the person or corporation to whom is addressed, and neither the report nor the name of these laboratories nor any members of its staff may be used in connection with the advertising, sale or any product or process, or any non-regulatory third party without the written permission of Hy-Laboratories Ltd. The results are approved for samples tested only. For research use only.

| Locus designation | 010HikeshiFbp2 | BGU 010iHikeshin4 p7 | 009HikeshiCTRFb p2 | BGU 009iHikeshiCTRn2p8 | 001STXBP1Fb p1 | BGU 001iSTXBP1n6p13 |
|---|---|---|---|---|---|---|
| D8S1179 | 10, 14 | 10, 14 | 8, 10 | 8, 10 | 10 | 10 |
| D21S11 | 31.2, 32 | 31.2, 32 | 30, 31.2 | 30, 31.2 | 28 | 28 |
| D7S820 | 11 | 11 | 10, 11 | 10,11 | 8, 11 | 8, 11 |
| CSF1PO | 8, 10 | 8, 10 | 8, 13 | 8, 13 | 12 | 12 |
| D3S1358 | 14, 16 | 14, 16 | 14, 17 | 14, 17 | 14, 18 | 14, 18 |
| TH01 | 6, 9.3 | 6, 9.3 | 9.3 | 9.3 | 6, 9 | 6, 9 |
| D13S317 | 8 | 8 | 8, 12 | 8, 12 | 12, 13 | 12, 13 |
| D16S539 | 11, 12 | 11, 12 | 8, 12 | 8, 12 | 12 | 12 |
| D2S1338 | 16, 25 | 16, 25 | 16, 23 | 16, 23 | 17, 24 | 17, 24 |
| D19S433 | 12, 14 | 12, 14 | 12, 15 | 12, 15 | 13, 14 | 13, 14 |
| vWA | 14, 16 | 14, 16 | 14, 18 | 14, 18 | 15, 17 | 15, 17 |
| TPOX | 8, 9 | 8, 9 | 8, 11 | 8, 11 | 8, 9 | 8, 9 |
| D18S51 | 13, 14 | 13, 14 | 13, 15 | 13, 15 | 14, 16 | 14, 16 |
| AMEL | X | X | X | X | X, Y | X, Y |
| D5S818 | 11, 13 | 11, 13 | 11, 12 | 11, 12 | 12, 13 | 12, 13 |
| FGA | 20, 24 | 20, 24 | 21, 24 | 21, 24 | 20 | 20 |

This report is submitted for the exclusive use of the person or corporation to whom is addressed, and neither the report nor the name of these laboratories nor any members of its staff may be used in connection with the advertising, sale or any product or process, or any non-regulatory third party without the written permission of Hy-Laboratories Ltd. The results are approved for samples tested only. For research use only.

| Locus designation | 002STX BP1CTRFbp2 | BGU 002iSTX BP1CTRn2p9 | 011PTHSFbp3 | BGU 011iPTHSn3p21 | 011PTHSCTRFbp 2 | BGU 011iPTHSCTRn5p21 |
|---|---|---|---|---|---|---|
| D8S1179 | 10, 14 | 10, 14 | 11, 13 | 11, 13 | 11, 15 | 11, 15 |
| D21S11 | 28, 32.2 | 28, 32.2 | 32.2 | 32.2 | 28, 32.2 | 28, 32.2 |
| D7S820 | 9, 12 | 9, 12 | 8, 11 | 8, 11 | 8, 11 | 8, 11 |
| CSF1PO | 12 | 12 | 10, 11 | 10, 11 | 6, 10 | 6, 10 |
| D3S1358 | 14, 17 | 14, 17 | 17 | 17 | 15, 17 | 15, 17 |
| TH01 | 8, 9 | 8, 9 | 7 | 7 | 7 | 7 |
| D13S317 | 12, 13 | 12,13 | 10, 12 | 10, 12 | 11, 12 | 11, 12 |
| D16S539 | 11 | 11 | 9, 11 | 9, 11 | 11, 13 | 11, 13 |
| D2S1338 | 19, 21 | 19, 21 | 18, 20 | 18, 20 | 17, 20 | 17, 20 |
| D19S433 | 13, 15 | 13, 15 | 13 | 13 | 13 | 13 |
| vWA | 15 | 15 | 18, 19 | 18, 19 | 16, 18 | 16, 18 |
| TPOX | 8, 9 | 8, 9 | 8, 12 | 8, 12 | 8, 12 | 8, 12 |
| D18S51 | 16, 17 | 16, 17 | 18, 19 | 18, 19 | 18, 19 | 18, 19 |
| AMEL | X, Y | X, Y | X, Y | X, Y | X, Y | X, Y |
| D5S818 | 12, 13 | 12, 13 | 11, 13 | 11, 13 | 11 | 11 |
| FGA | 20, 22 | 20, 22 | 22, 26 | 22, 26 | 26 | 26 |

This report is submitted for the exclusive use of the person or corporation to whom is addressed, and neither the report nor the name of these laboratories nor any members of its staff may be used in connection with the advertising, sale or any product or process, or any non-regulatory third party without the written permission of Hy-Laboratories Ltd. The results are approved for samples tested only. For research use only.

| Locus designation | 012PTHSCTRFbp3 | BGU 012iPTHSCTRn6p15 | 013PTHSFbp2 | BGU 013iPTHSn3p20 | 013 PTHSCTRFbp3 | BGU 013iPTHSCTRn4p22 |
|---|---|---|---|---|---|---|
| D8S1179 | 11, 13 | 11, 13 | 13, 14 | 13, 14 | 13, 15 | 13, 15 |
| D21S11 | 26, 31 | 26, 31 | 29, 30 | 29, 30 | 29, 30 | 29, 30 |
| D7S820 | 8, 11 | 8, 11 | 9, 10 | 9, 10 | 9, 10 | 9, 10 |
| CSF1PO | 10, 12 | -- | 11, 12 | 11, 12 | 10, 12 | 10, 12 |
| D3S1358 | 16, 17 | 16, 17 | 15, 18 | 15, 18 | 15, 17 | 15, 17 |
| TH01 | 9.3 | 9.3 | 9, 9.3 | 9, 9.3 | 8, 9.3 | 8, 9.3 |
| D13S317 | 13, 14 | 13, 14 | 8, 14 | 8, 14 | 8 | 8 |
| D16S539 | 13 | 13 | 10, 12 | 10, 12 | 11, 12 | 11, 12 |
| D2S1338 | 20, 25 | -- | 17, 21 | 17, 21 | 17, 21 | 17, 21 |
| D19S433 | 13, 14 | 13, 14 | 14, 14.2 | 14, 14.2 | 14, 14.2 | 14, 14.2 |
| vWA | 14, 19 | 14, 19 | 16, 17 | 16, 17 | 16, 19 | 16, 19 |
| TPOX | 8, 10 | 8, 10 | 9 | 9 | 8, 9 | 8, 9 |
| D18S51 | 13, 17 | 13, 17 | 12, 13 | 12, 13 | 13, 14 | 13, 14 |
| AMEL | X | X | X, Y | X, Y | X, Y | X, Y |
| D5S818 | 12, 13 | 12, 13 | 9 | 9 | 9, 10 | 9, 10 |
| FGA | 23, 24 | 23, 24 | 23, 24 | 23, 24 | 23 | 23 |

Performed by: **Sveta Gaiduk** **(Name & Sign)** DATE: **25.08.2021**

Reviewed by: **Dr. Ortal Shimon** **(Name & Sign)** DATE: **25.08.2021**

This report is submitted for the exclusive use of the person or corporation to whom is addressed, and neither the report nor the name of these laboratories nor any members of its staff may be used in connection with the advertising, sale or any product or process, or any non-regulatory third party without the written permission of Hy-Laboratories Ltd. The results are approved for samples tested only. For research use only.
