## Supplementary material for "Generation and characterization of iPSC lines (UOHi001-A, UOHi002-A) From a patient with SHANK3 mutation and her healthy mother": Supp 2

| Sample File | Sample Name | Panel | OS | SQ |
|---|---|---|---|---|
| STR_G5_run395825a_A01_BGU009iHikeshin1p7.fsa | BGU009iHikeshin1p7 | IF_3730_GM_Panel_CLA_v1 | | |

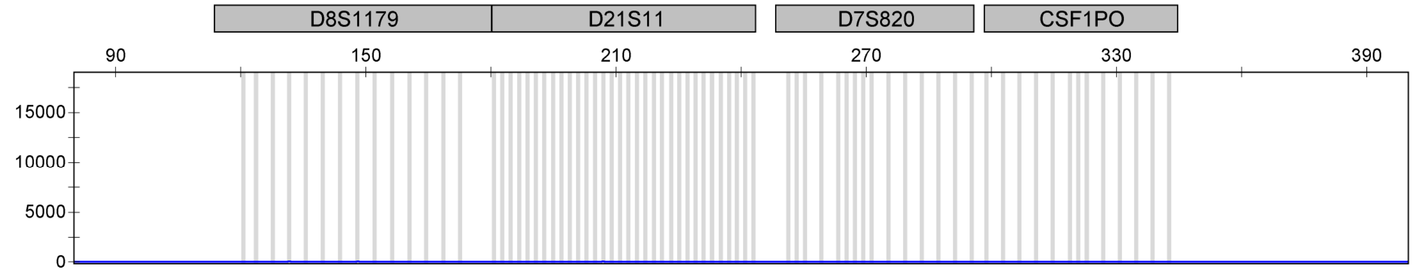

| | | |
|---|---|---|
| STR_G5_run395825a_A01_BGU009iHikeshin1p7.fsa | BGU009iHikeshin1p7 | IF_3730_GM_Panel_CLA_v1 |

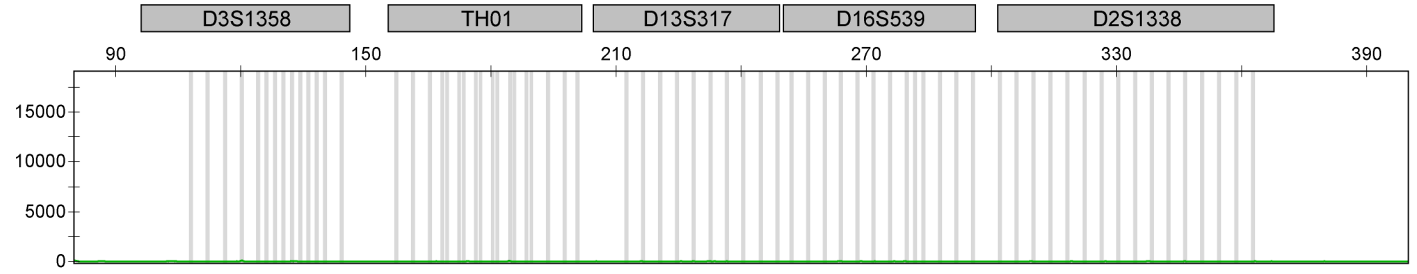

| | | |
|---|---|---|
| STR_G5_run395825a_A01_BGU009iHikeshin1p7.fsa | BGU009iHikeshin1p7 | IF_3730_GM_Panel_CLA_v1 |

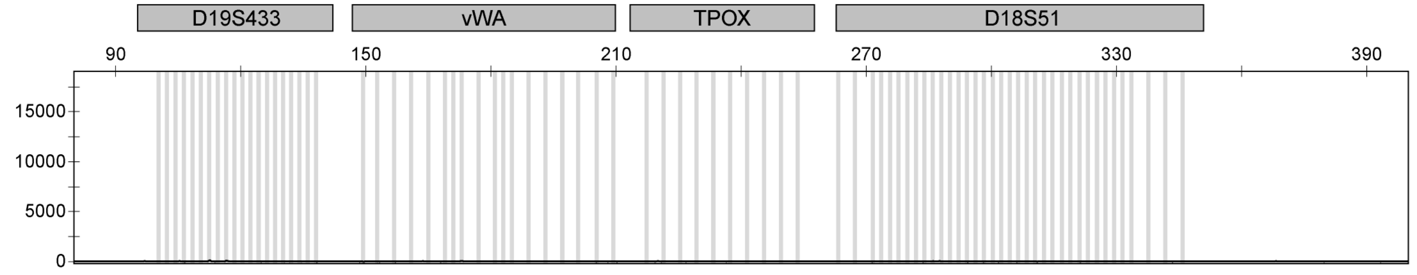

| | | |
|---|---|---|
| STR_G5_run395825a_A01_BGU009iHikeshin1p7.fsa | BGU009iHikeshin1p7 | IF_3730_GM_Panel_CLA_v1 |

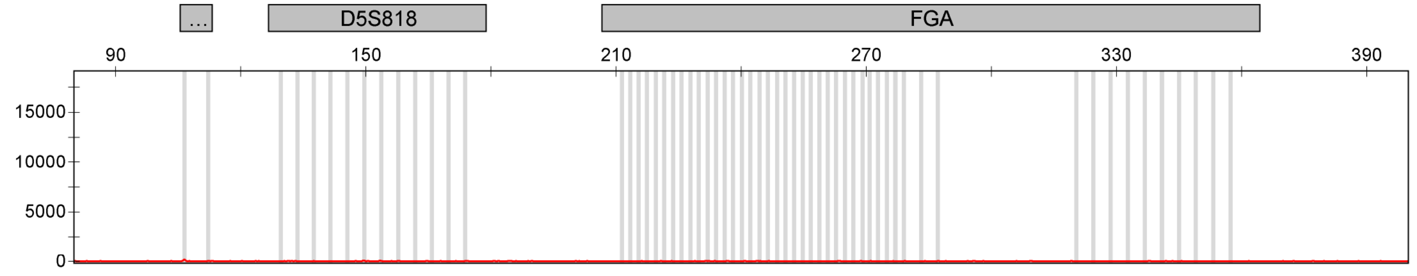

| Sample File | Sample Name | Panel | OS | SQ |
|---|---|---|---|---|
| STR_G5_run395825a_B01_009HikeshiCTRFbp2.fsa | 009HikeshiCTRFbp2 | IF_3730_GM_Panel_CLA_v1 | | |

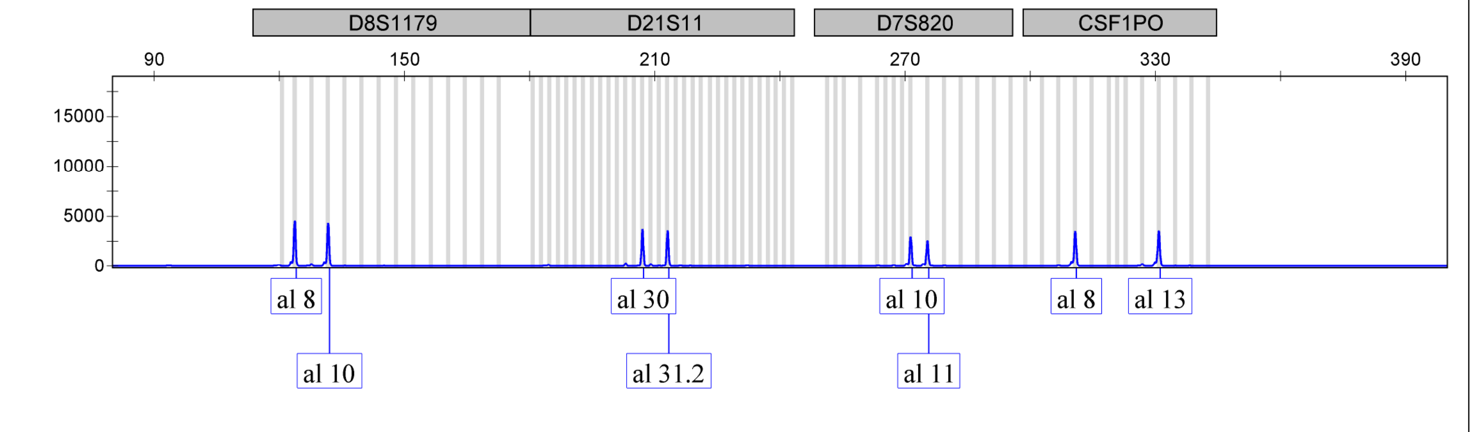

| | | |
|---|---|---|
| STR_G5_run395825a_B01_009HikeshiCTRFbp2.fsa | 009HikeshiCTRFbp2 | IF_3730_GM_Panel_CLA_v1 |

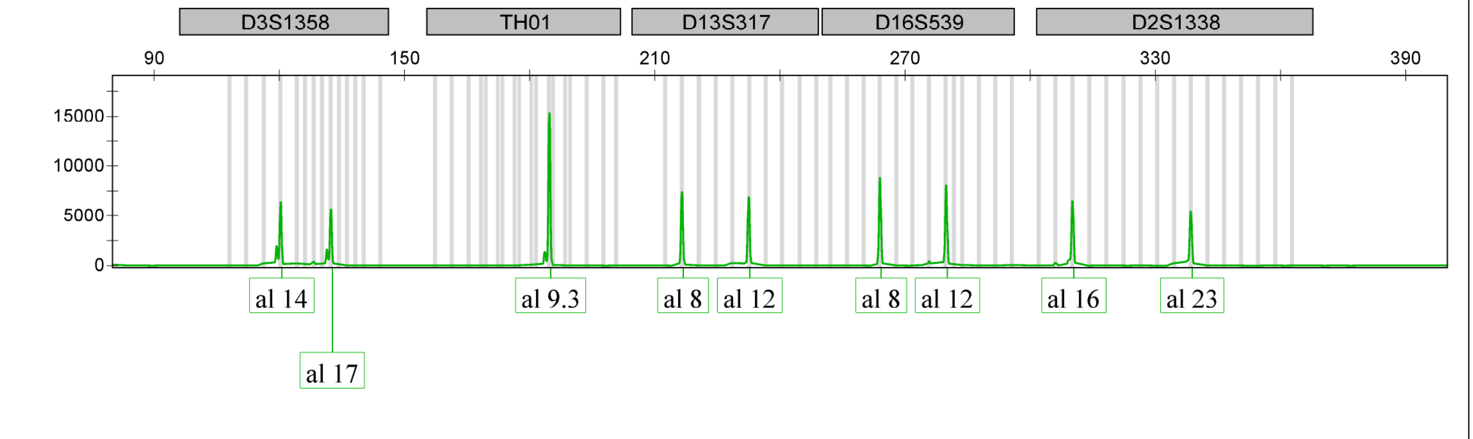

| | | |
|---|---|---|
| STR_G5_run395825a_B01_009HikeshiCTRFbp2.fsa | 009HikeshiCTRFbp2 | IF_3730_GM_Panel_CLA_v1 |

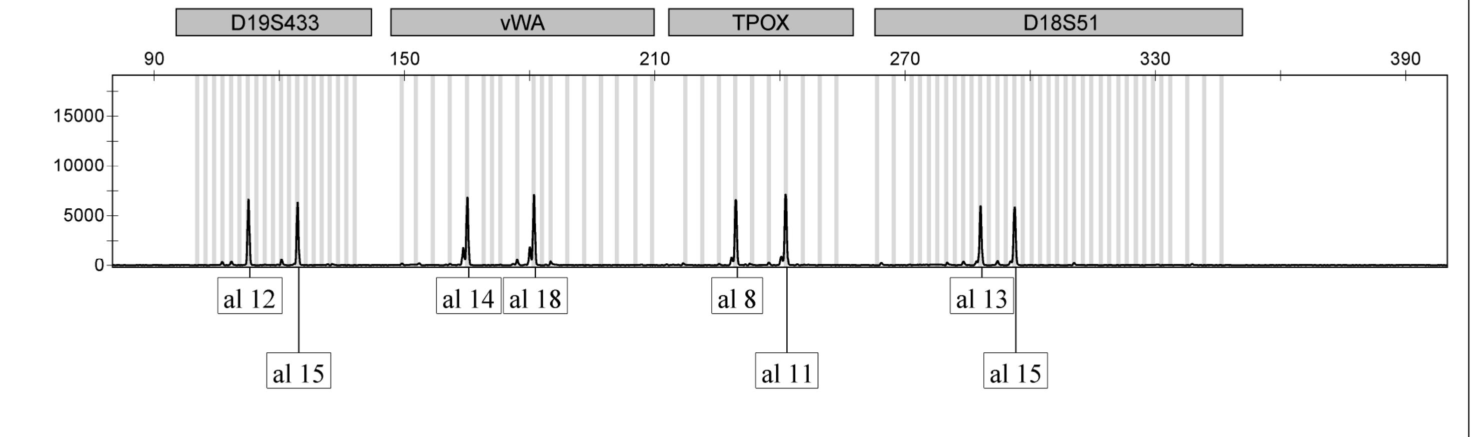

Applied Biosystems
GeneMapper Software 5

Reap_395825

| Sample File | Sample Name | Panel | OS | SQ |
|---|---|---|---|---|
| STR_G5_run395825a_B01_009HikeshiCTRFbp2.fsa | 009HikeshiCTRFbp2 | IF_3730_GM_Panel_CLA_v1 | | |

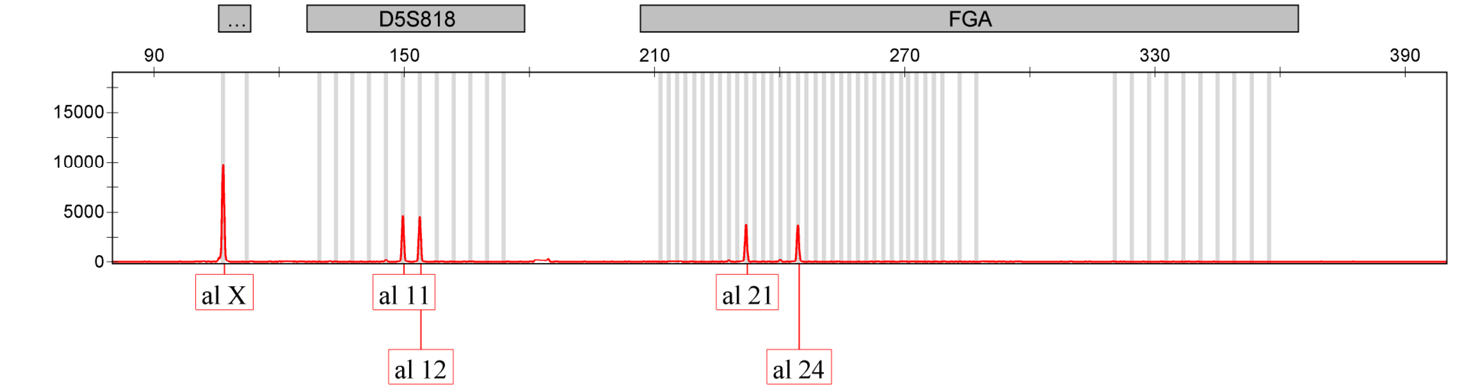

| | | |
|---|---|---|
| STR_G5_run395825a_C01_BGU011iPTHSn3p21.fsa | BGU011iPTHSn3p21 | IF_3730_GM_Panel_CLA_v1 |

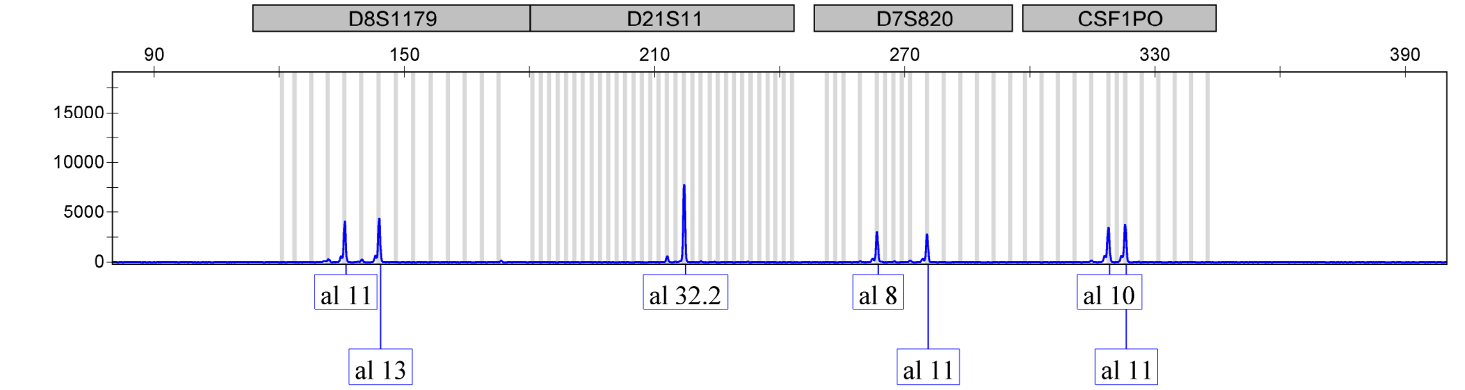

| | | |
|---|---|---|
| STR_G5_run395825a_C01_BGU011iPTHSn3p21.fsa | BGU011iPTHSn3p21 | IF_3730_GM_Panel_CLA_v1 |

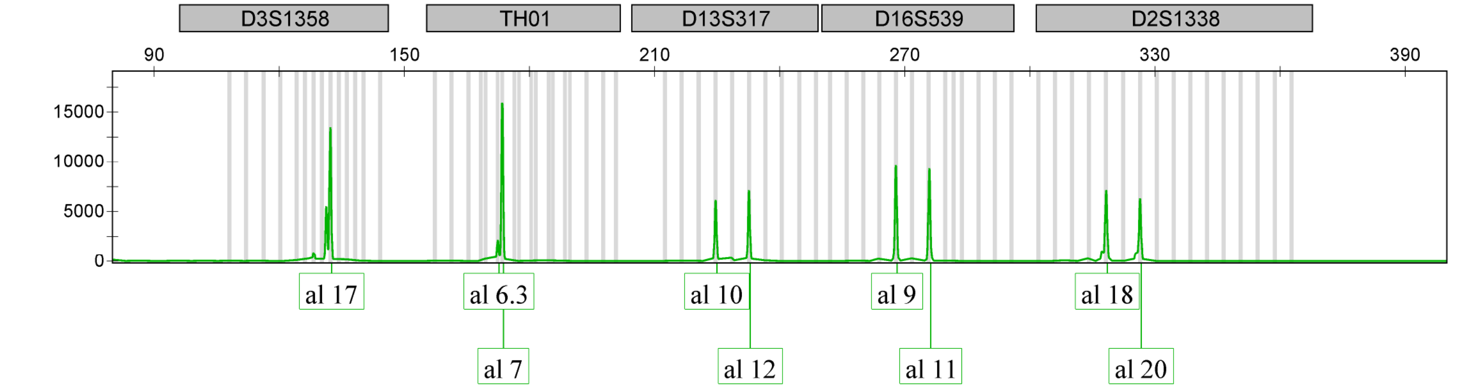

| Sample File | Sample Name | Panel | OS | SQ |
|---|---|---|---|---|
| STR_G5_run395825a_C01_BGU011iPTHSn3p21.fsa | BGU011iPTHSn3p21 | IF_3730_GM_Panel_CLA_v1 | | |

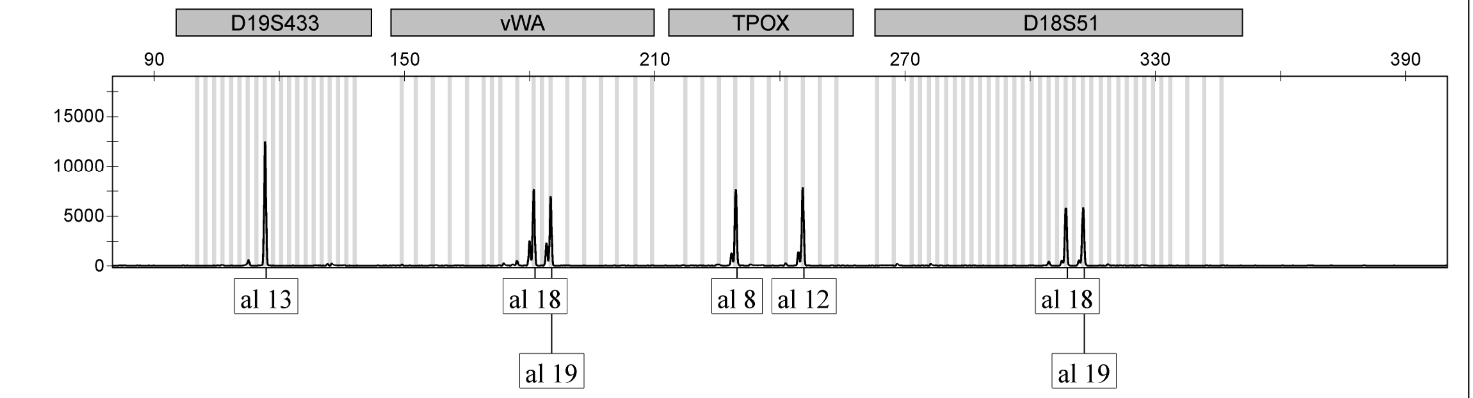

| | | |
|---|---|---|
| STR_G5_run395825a_C01_BGU011iPTHSn3p21.fsa | BGU011iPTHSn3p21 | IF_3730_GM_Panel_CLA_v1 |

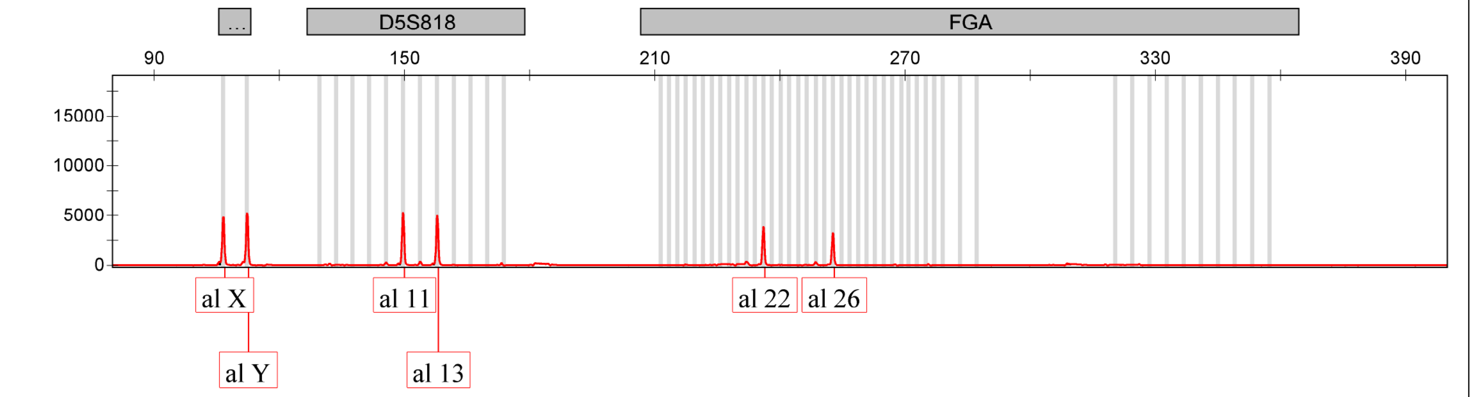

| | | |
|---|---|---|
| STR_G5_run395825a_D01_BGU012iPTHSCTRn6p15.fsa | BGU012iPTHSCTRn6p15 | IF_3730_GM_Panel_CLA_v1 |

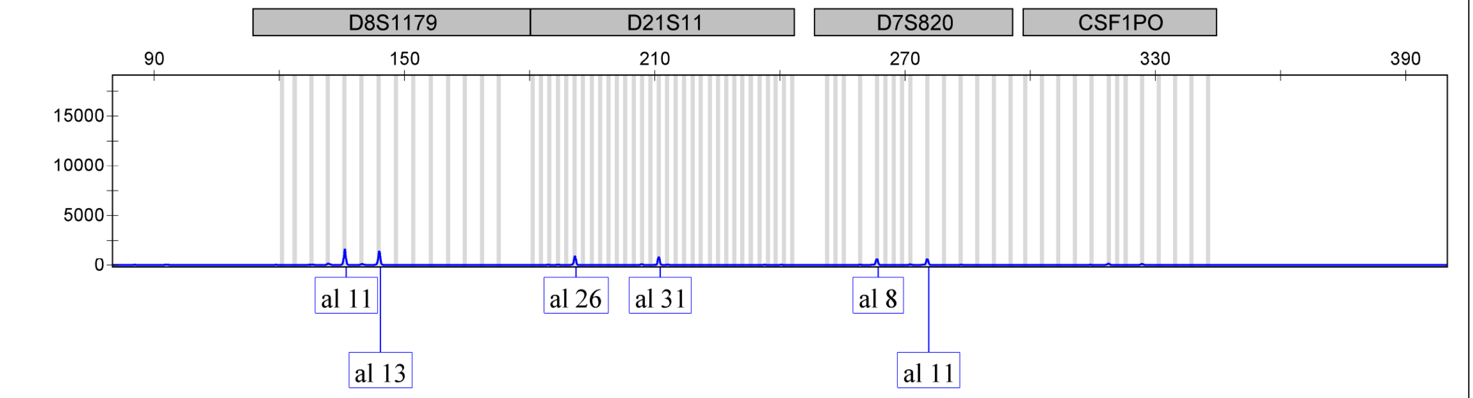

| Sample File | Sample Name | Panel | OS | SQ |
|---|---|---|---|---|
| STR_G5_run395825a_D01_BGU012iPTHSCTRn6p15.fsa | BGU012iPTHSCTRn6p15 | IF_3730_GM_Panel_CLA_v1 | | |

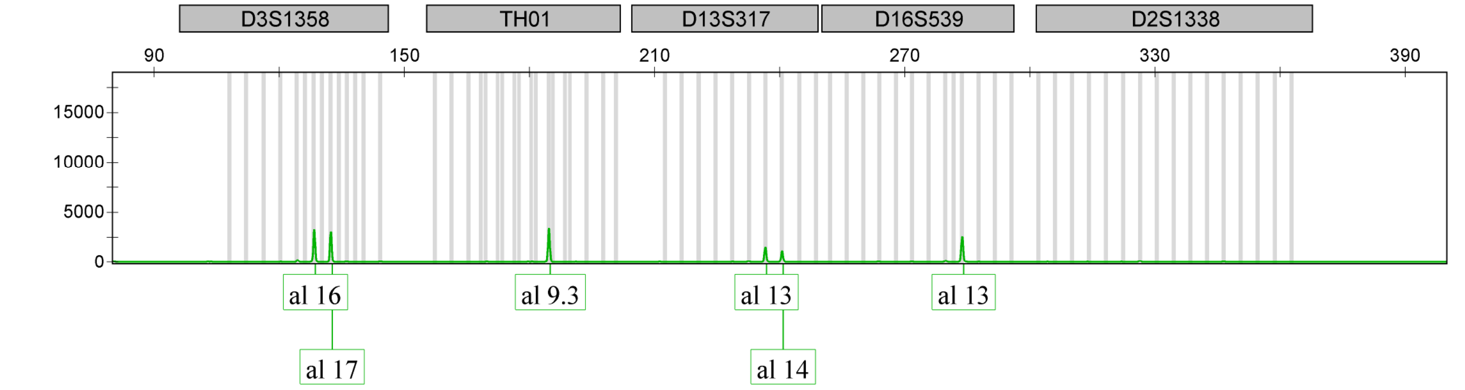

| STR_G5_run395825a_D01_BGU012iPTHSCTRn6p15.fsa | BGU012iPTHSCTRn6p15 | IF_3730_GM_Panel_CLA_v1 |
|---|---|---|

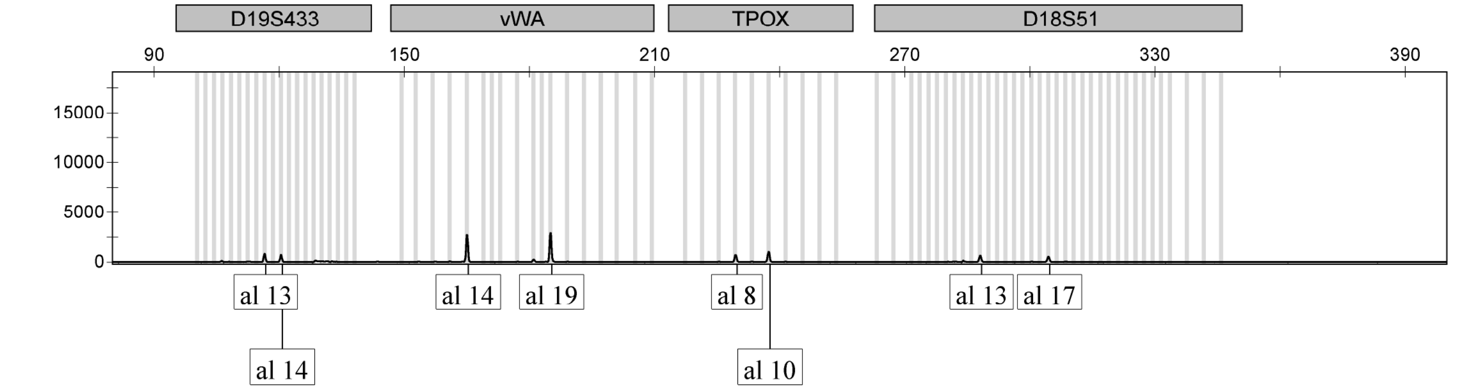

| STR_G5_run395825a_D01_BGU012iPTHSCTRn6p15.fsa | BGU012iPTHSCTRn6p15 | IF_3730_GM_Panel_CLA_v1 |
|---|---|---|

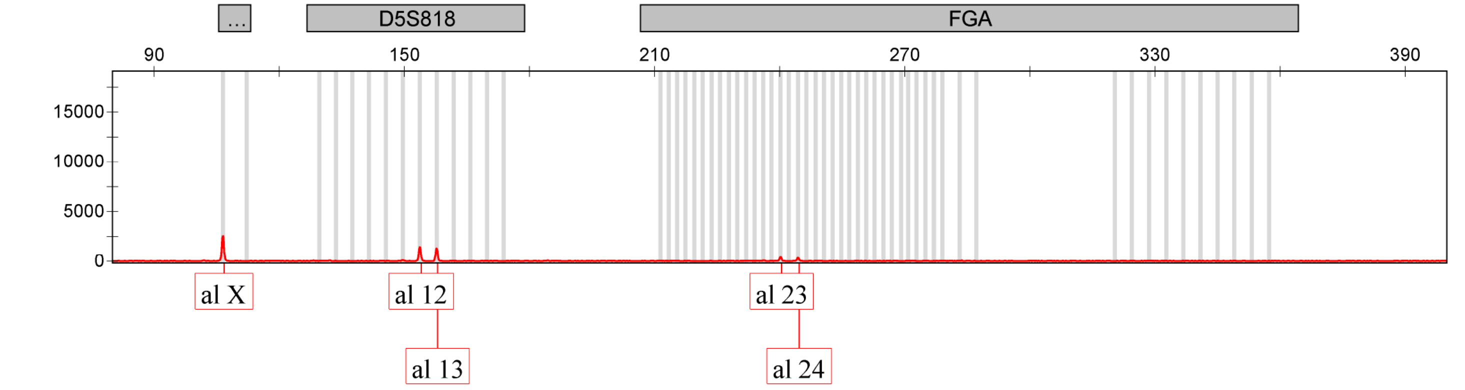

| Sample File | Sample Name | Panel | OS | SQ |
|---|---|---|---|---|
| STR_G5_run395825a_E01_pos.fsa | pos | IF_3730_GM_Panel_CLA_v1 | | |

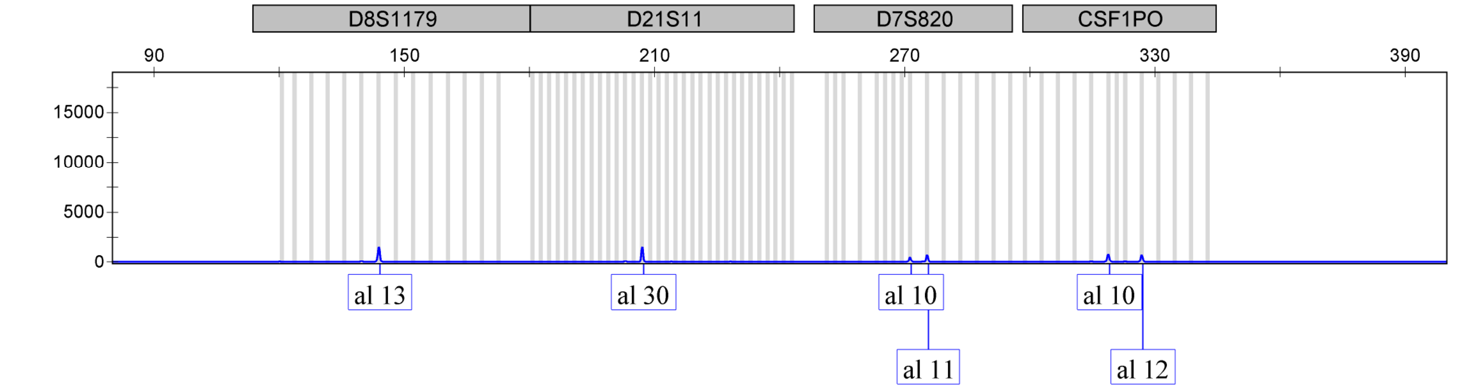

| | | |
|---|---|---|
| STR_G5_run395825a_E01_pos.fsa | pos | IF_3730_GM_Panel_CLA_v1 |

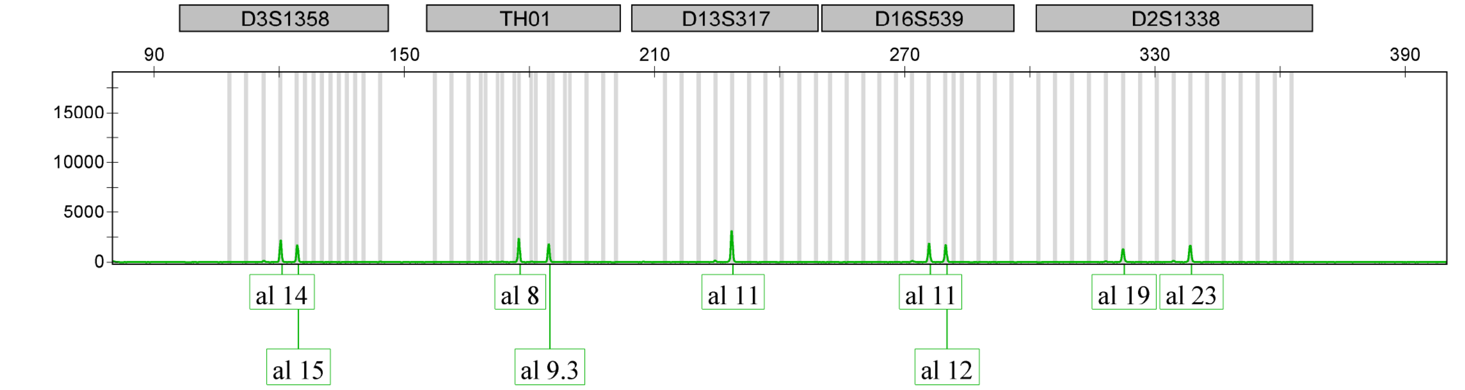

| | | |
|---|---|---|
| STR_G5_run395825a_E01_pos.fsa | pos | IF_3730_GM_Panel_CLA_v1 |

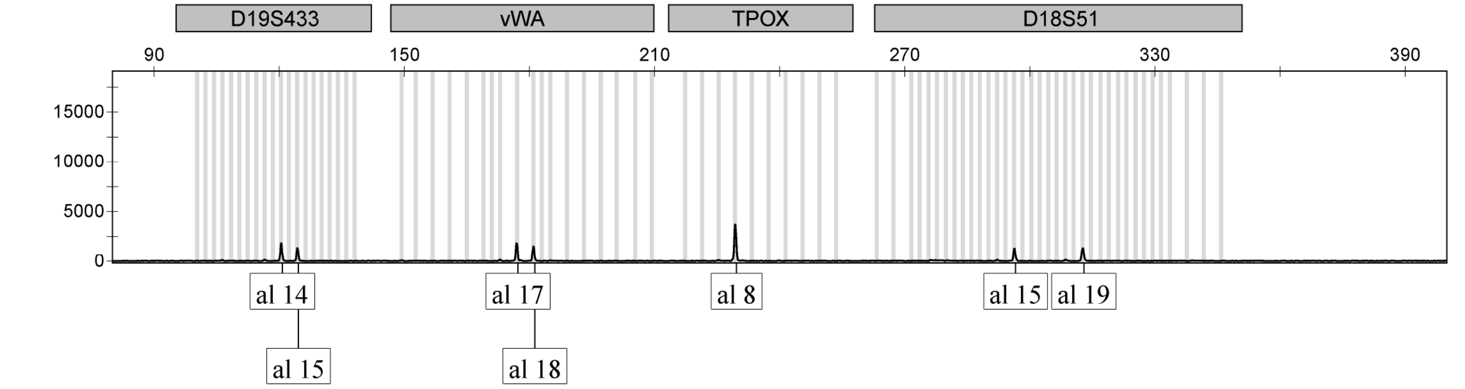

| Sample File | Sample Name | Panel | OS | SQ |
|---|---|---|---|---|
| STR_G5_run395825a_E01_pos.fsa | pos | IF_3730_GM_Panel_CLA_v1 | | |

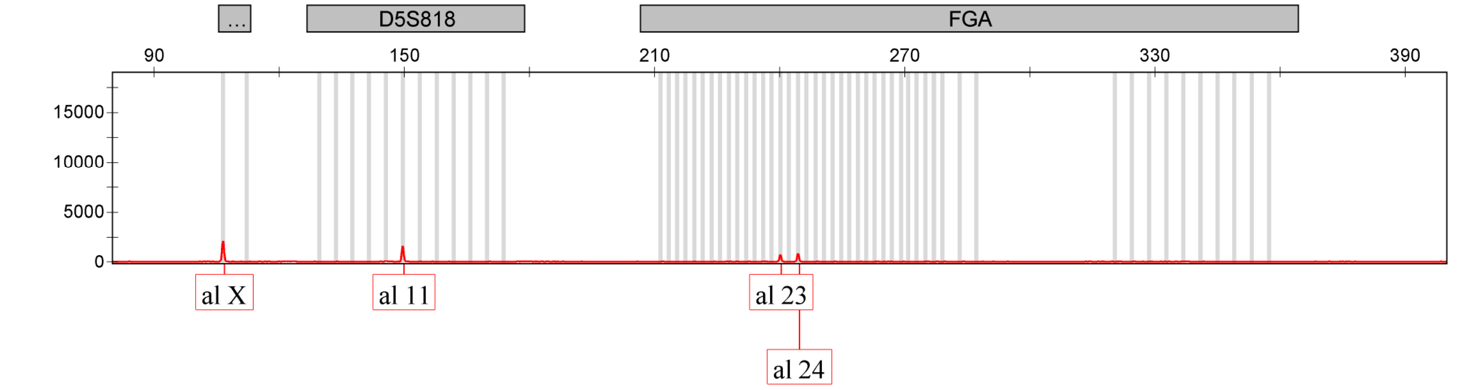

| | | |
|---|---|---|
| STR_G5_run395825a_F01_ladder.fsa | ladder | IF_3730_GM_Panel_CLA_v1 |

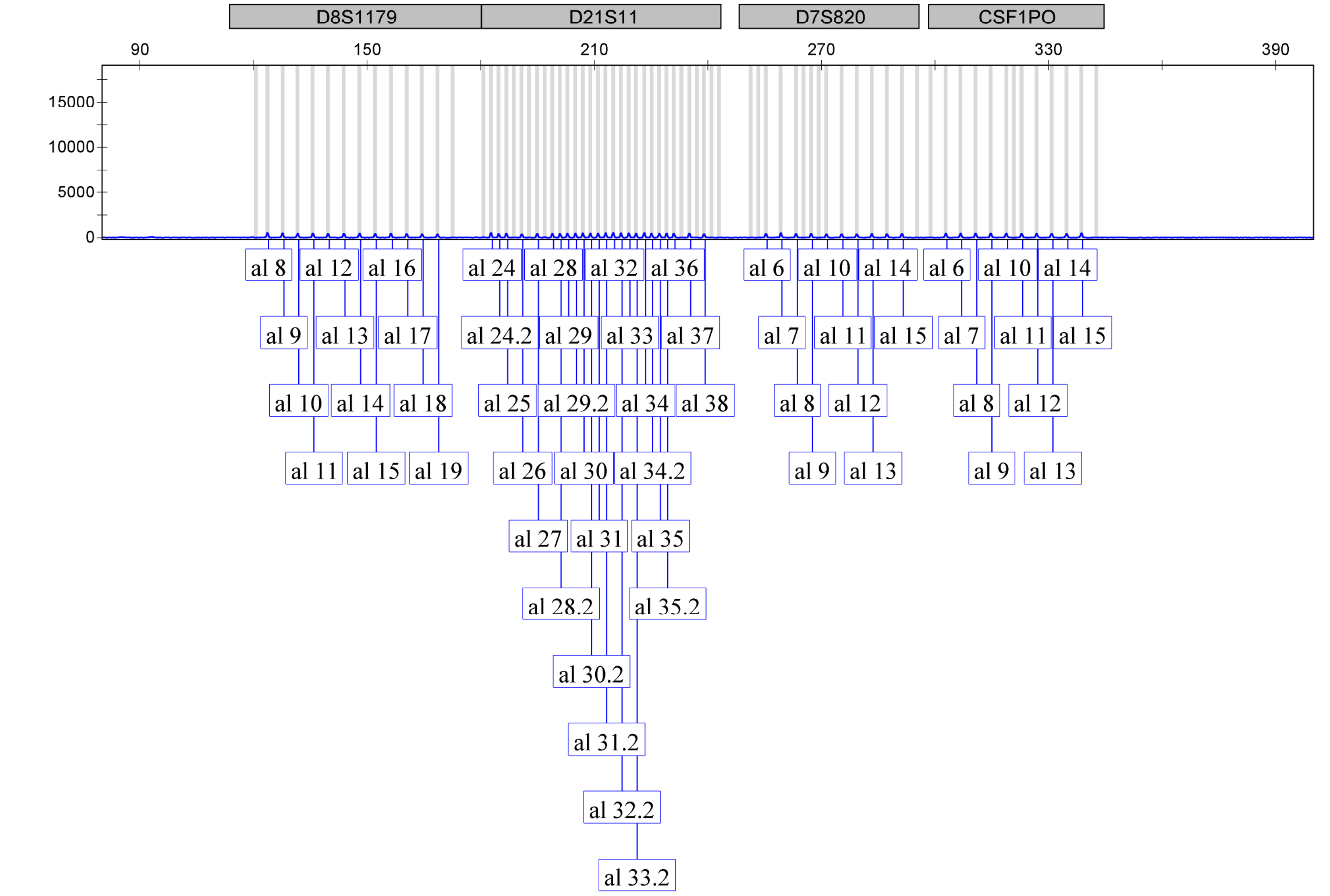

Applied Biosystems
GeneMapper Software 5

Reap_395825

| Sample File | Sample Name | Panel | OS | SQ |
|---|---|---|---|---|
| STR_G5_run395825a_F01_ladder.fsa | ladder | IF_3730_GM_Panel_CLA_v1 | | |

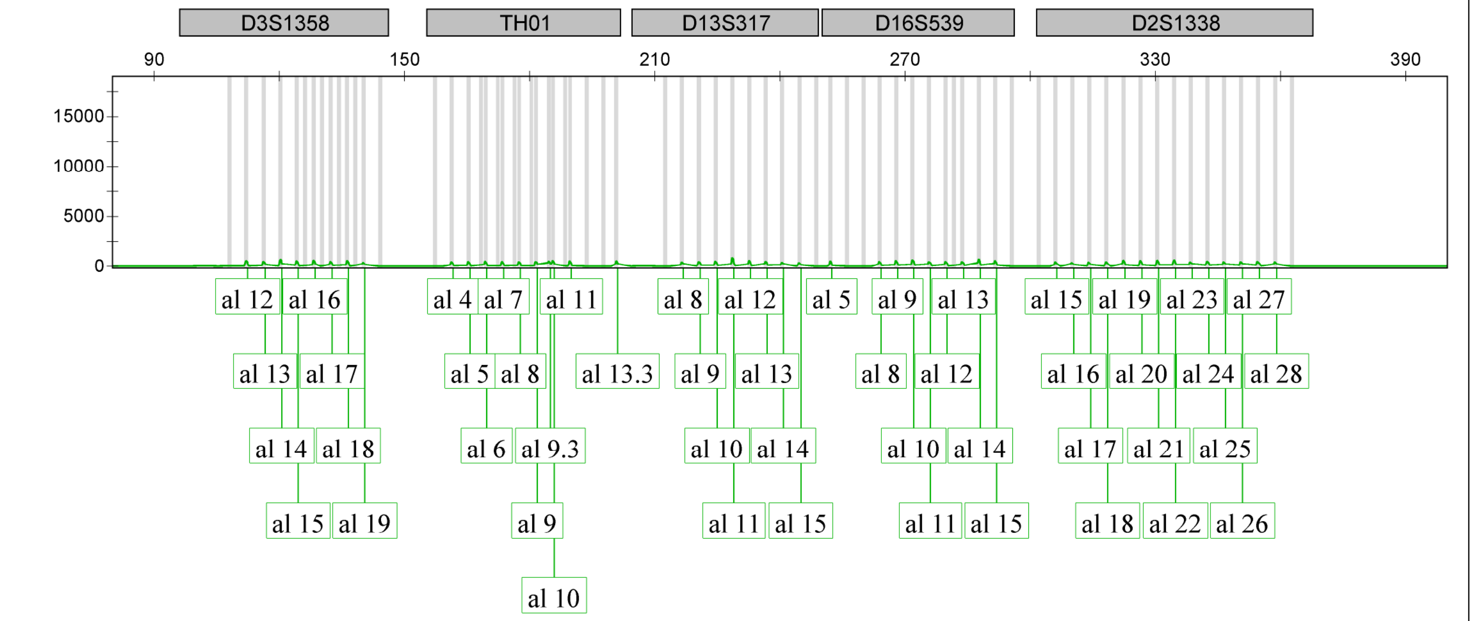

| Sample File | Sample Name | Panel | OS | SQ |
|---|---|---|---|---|
| STR_G5_run395825a_F01_ladder.fsa | ladder | IF_3730_GM_Panel_CLA_v1 | | |

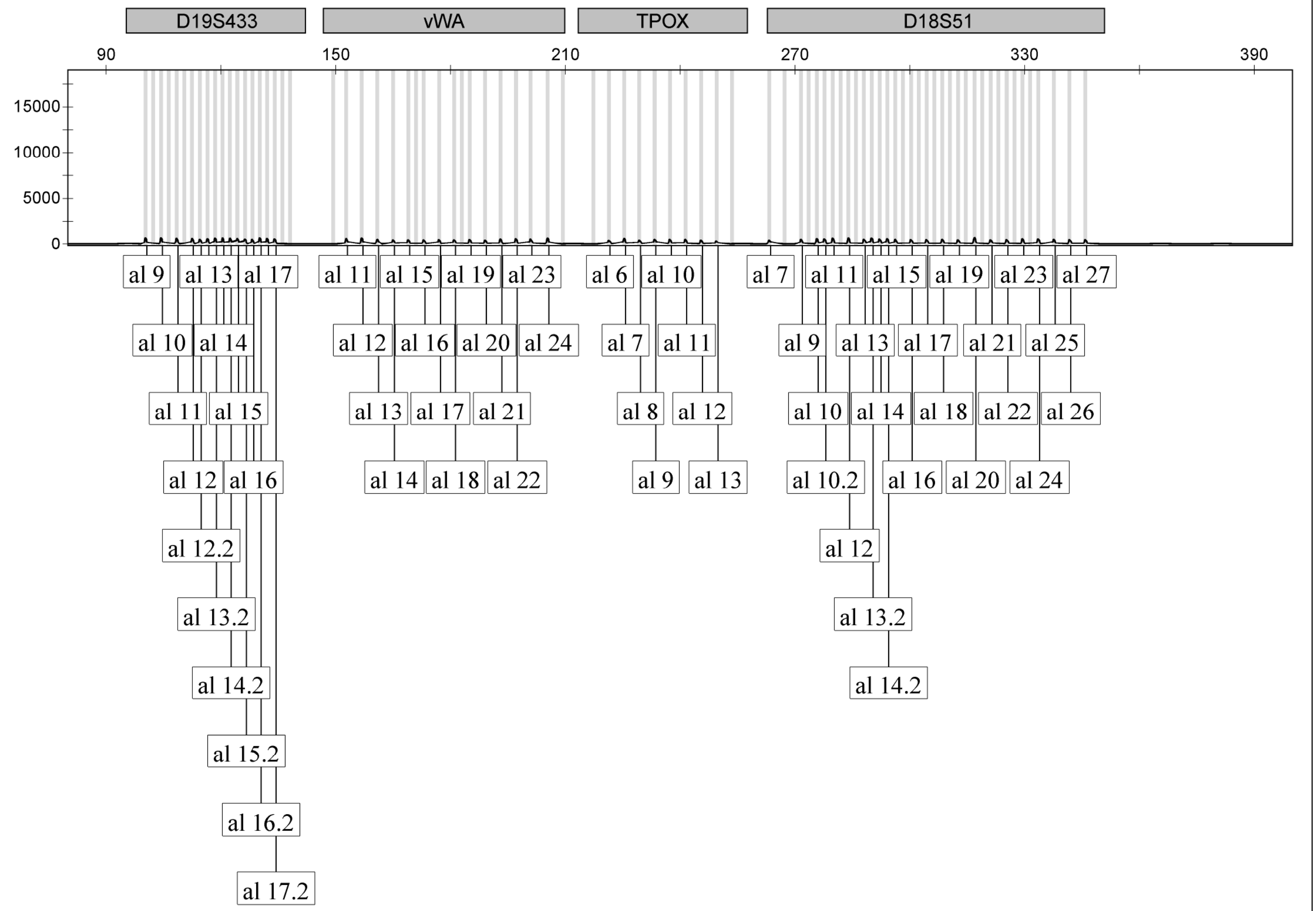

| Sample File | Sample Name | Panel | OS | SQ |
|---|---|---|---|---|
| STR_G5_run395825a_F01_ladder.fsa | ladder | IF_3730_GM_Panel_CLA_v1 | | |

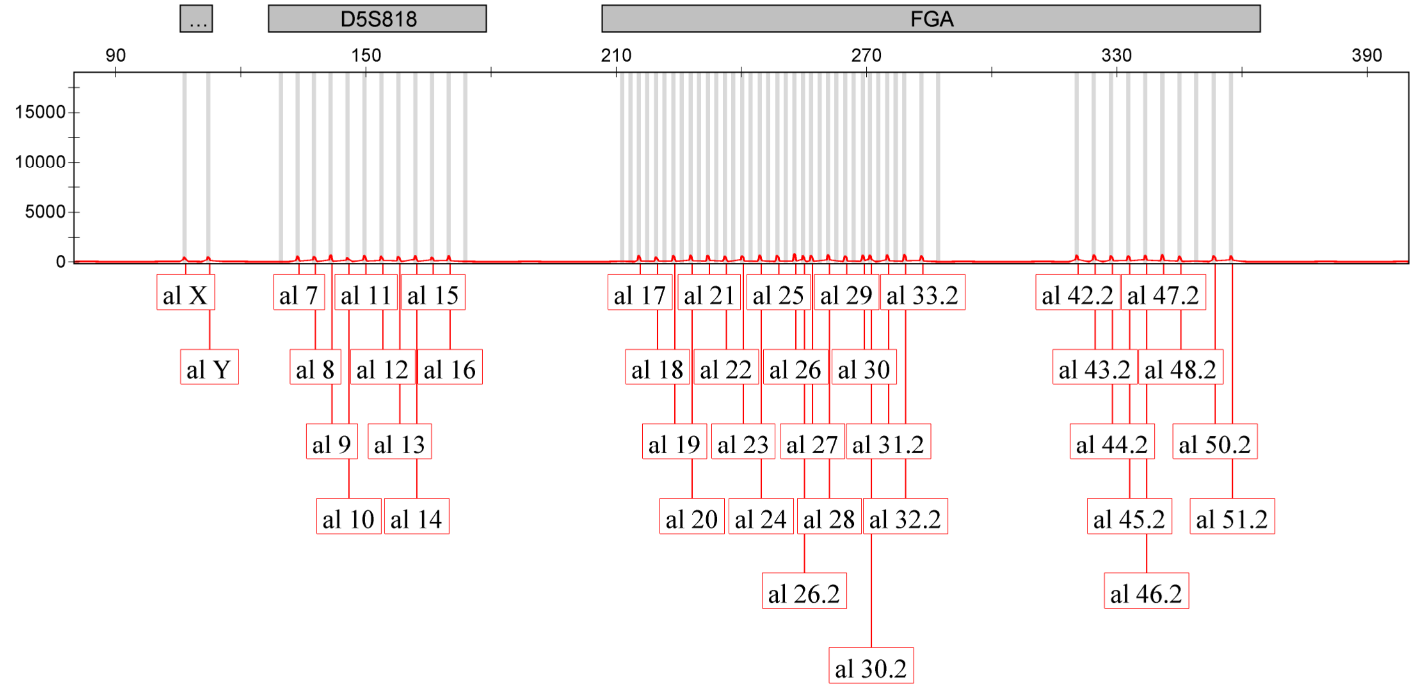
